## Supplementary Figures for "SVCurator: A Crowdsourcing app to visualize evidence of structural variants for the human genome"

**Supplementary Figure 1-5**

A)

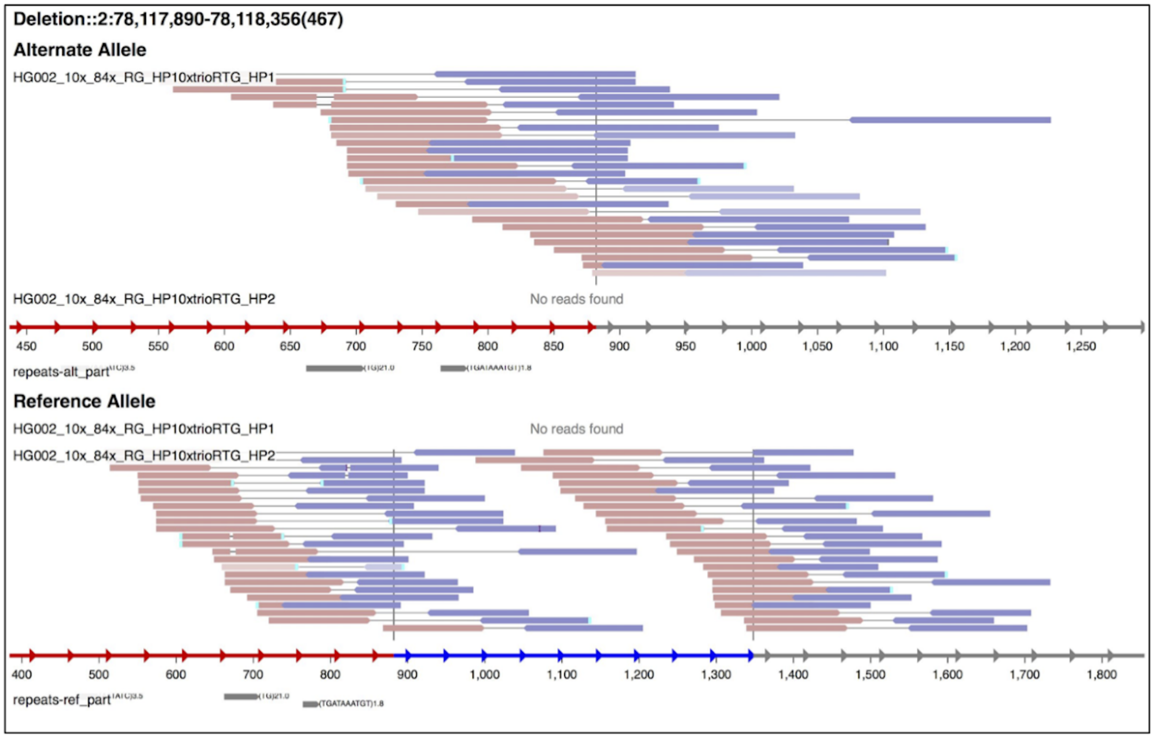


**Supplementary Figure 1**. Images were generated for each event from Integrated Genome Viewer, svviz2. A putative 467bp deletion is shown. Svviz2 generates read aligned images for each short read and long read sequencing technology. A) svviz2 read aligned image - 10x Genomics (read length = 98bp; read depth = 50x). Reads were aligned to reference and alternate allele by svviz2.

B)

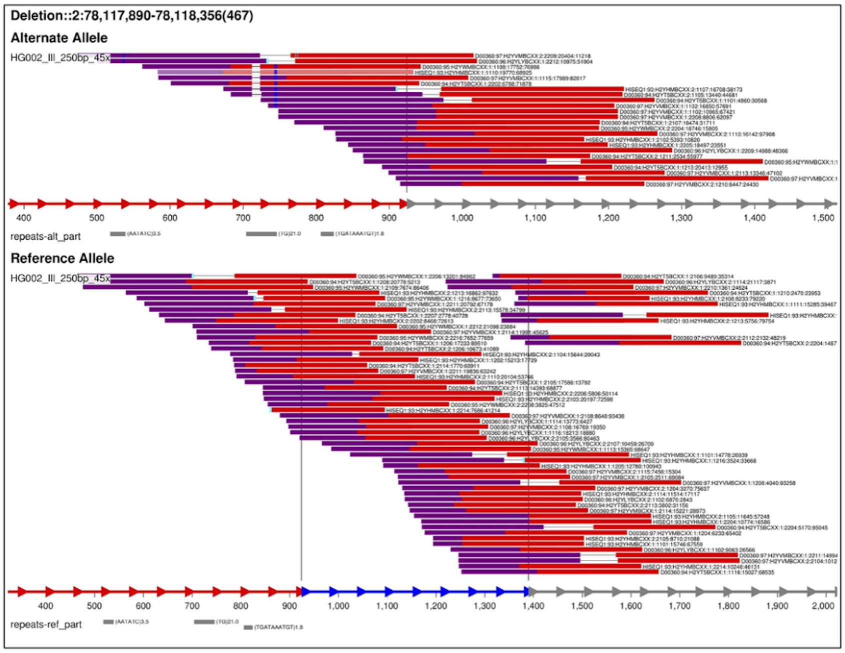


B) svviz2 read aligned image - Illumina HiSeq (read length = 250 bps; read depth = 40-50x)

C)

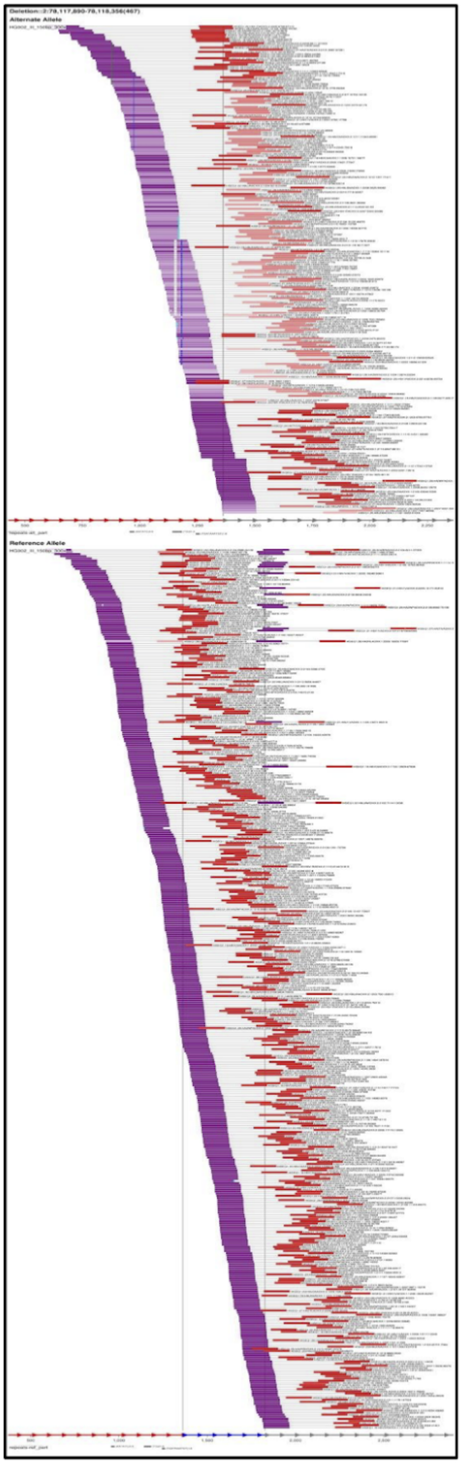

C) svviz2 read aligned image - Illumina HiSeq (read length = 148 bps; read depth = 296.83x)

D)

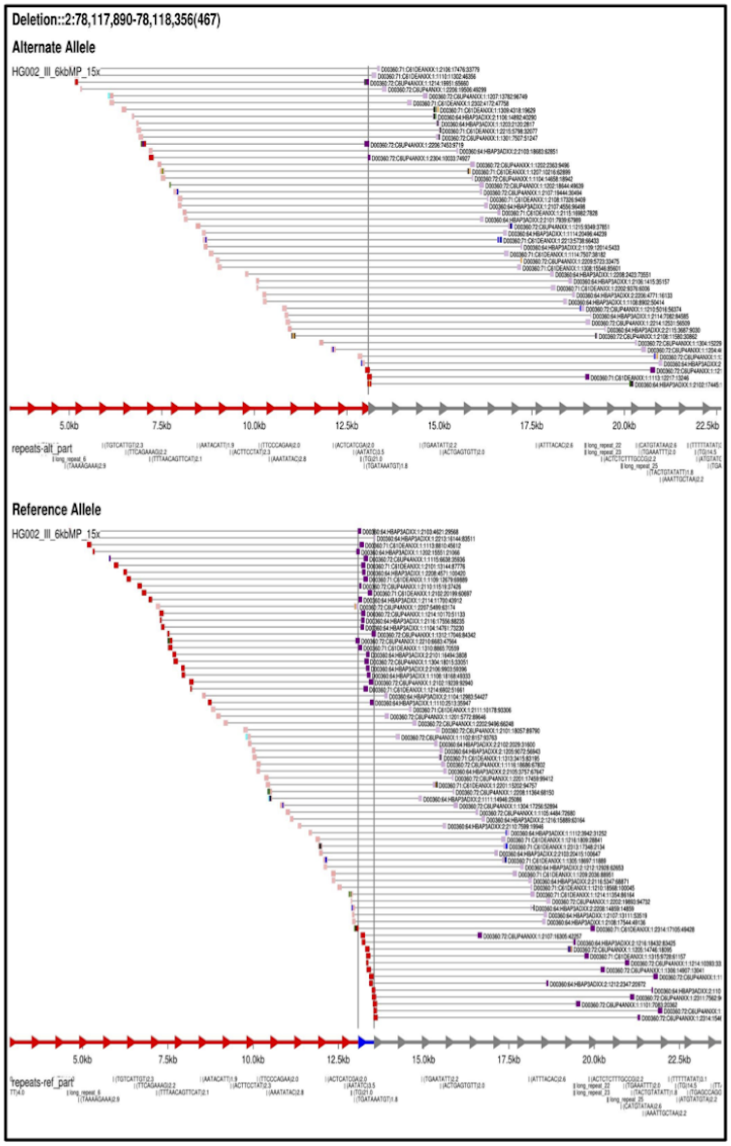


D) svviz2 read aligned image - Illumina Mate Pair (read length = 100 bps; insert size = 6000bp; read depth = 13-14x)

E)

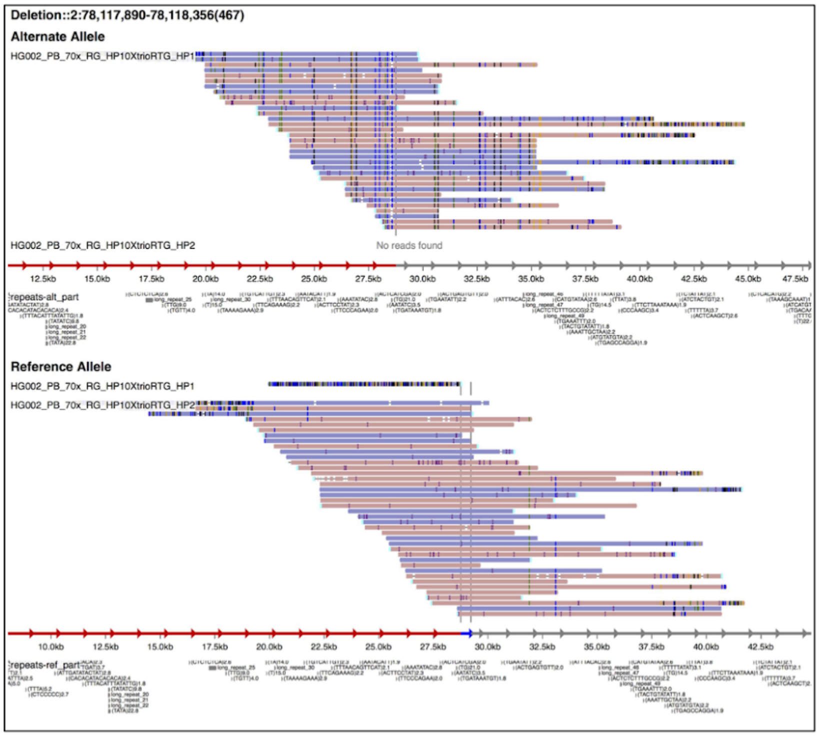


E) svviz2 read aligned image - Haplotype separated PacBio (read length = 10-11kb; read depth = 69x). Reads were haplotype separated using WhatsHap and aligned to reference and alternate allele by svviz2.

F)

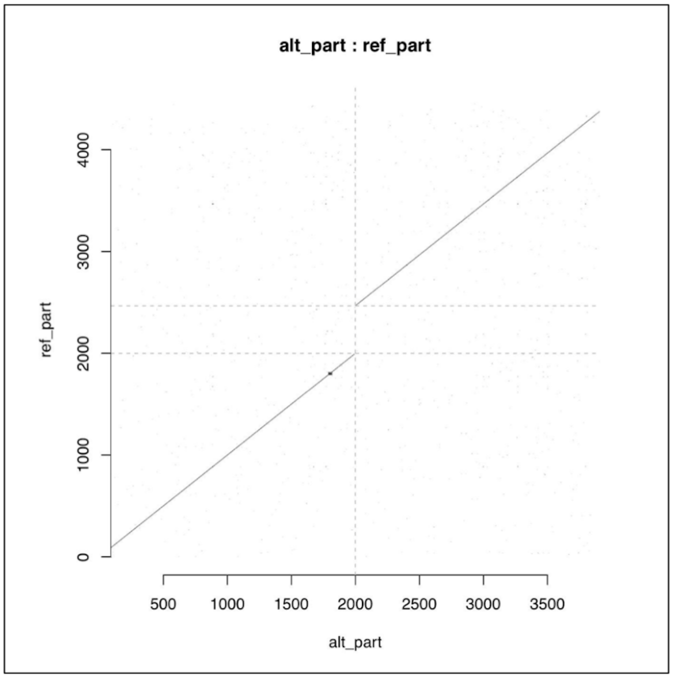


F) Svviz2 dotplot displaying reference versus alternate allele

G)

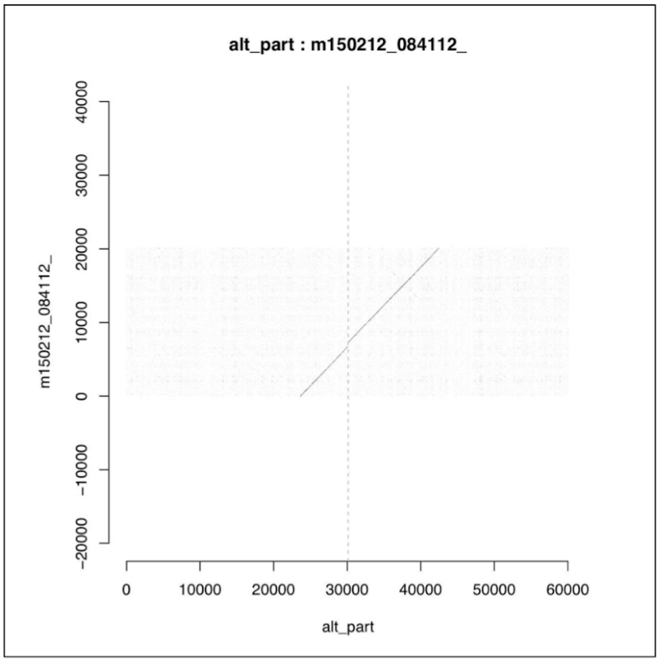


G) Svviz2 dotplot PacBio read with the highest mapping quality score versus the alternate allele.

H)

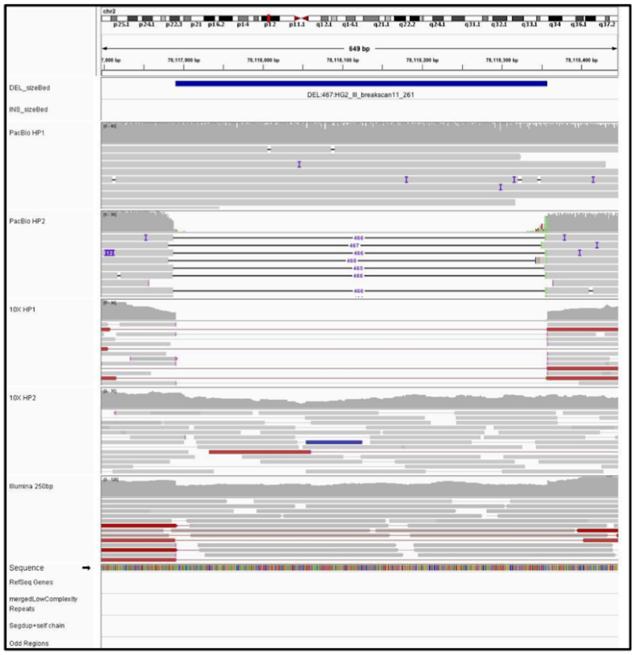


H) IGV image showing reads aligned to a putative variant. IGV tracks include: Haplotype separated PacBio reads, Haplotype Separated 10x Genomics reads, Illumina 250x250bp paired end sequencing, and tracks to describe repeat regions (low complexity repeats and segmental duplications). All reads were aligned to GRCh37 human reference genome.

I)

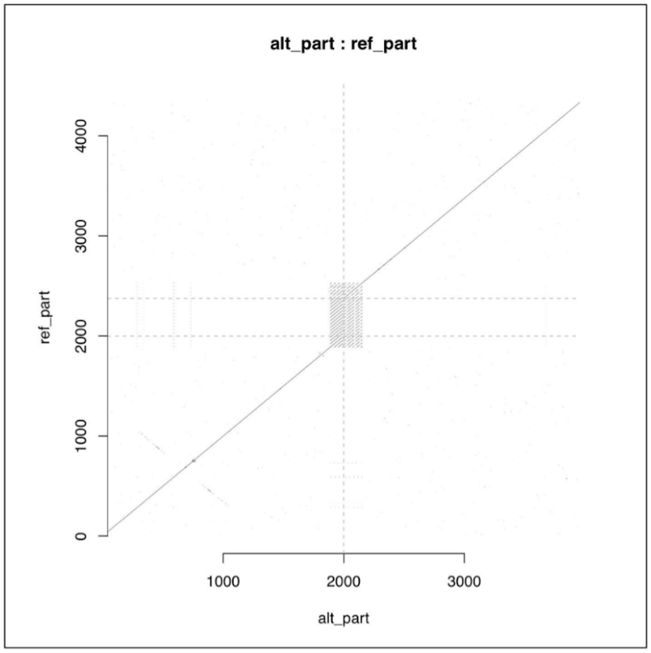


I) Svviz2 dotplot displaying a putative deletion in a tandem repeat.

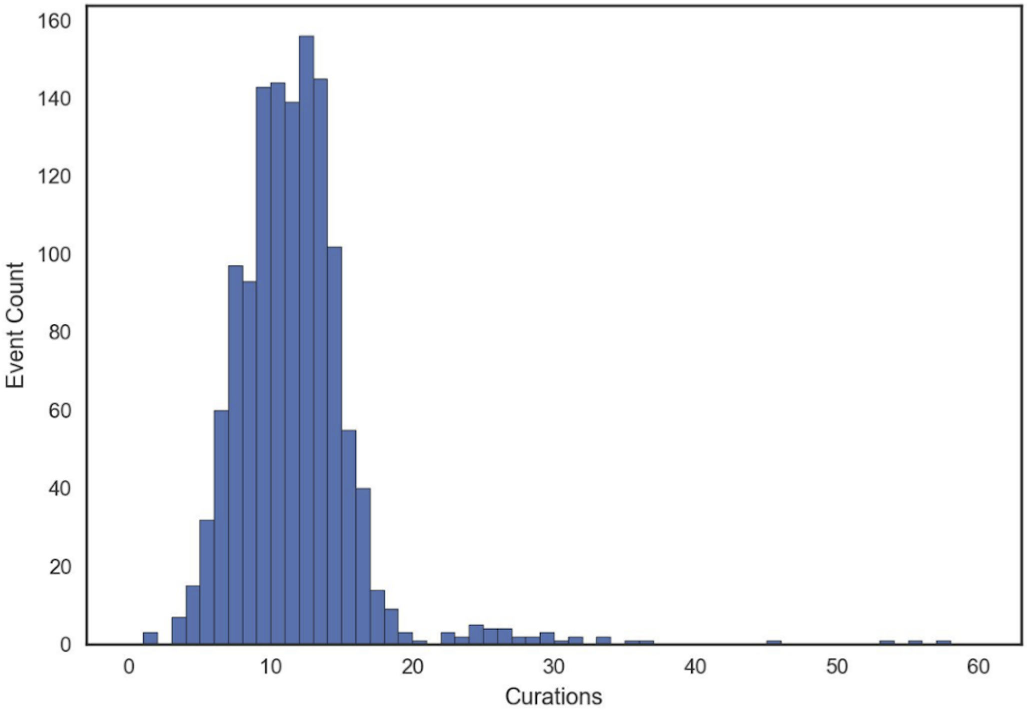


**Supplementary Figure 2.** Summary of the number of curations for each event. 61 curators evaluated SVCurator events. Each of the 1295 sites were curated on average 11 times with 1290 events curated at least 3 times.

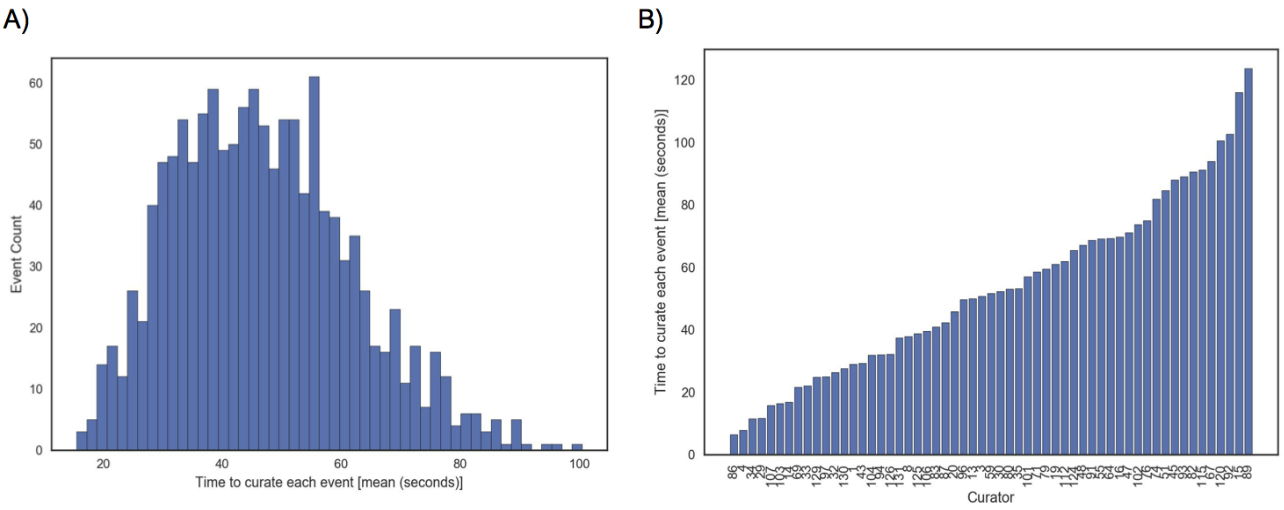


**Supplementary Figure 3.** An evaluation of the time to curate each SVCurator event. A) Overall distribution of the average time to curate events. B) Distribution of the average time to curate each event for each curator.

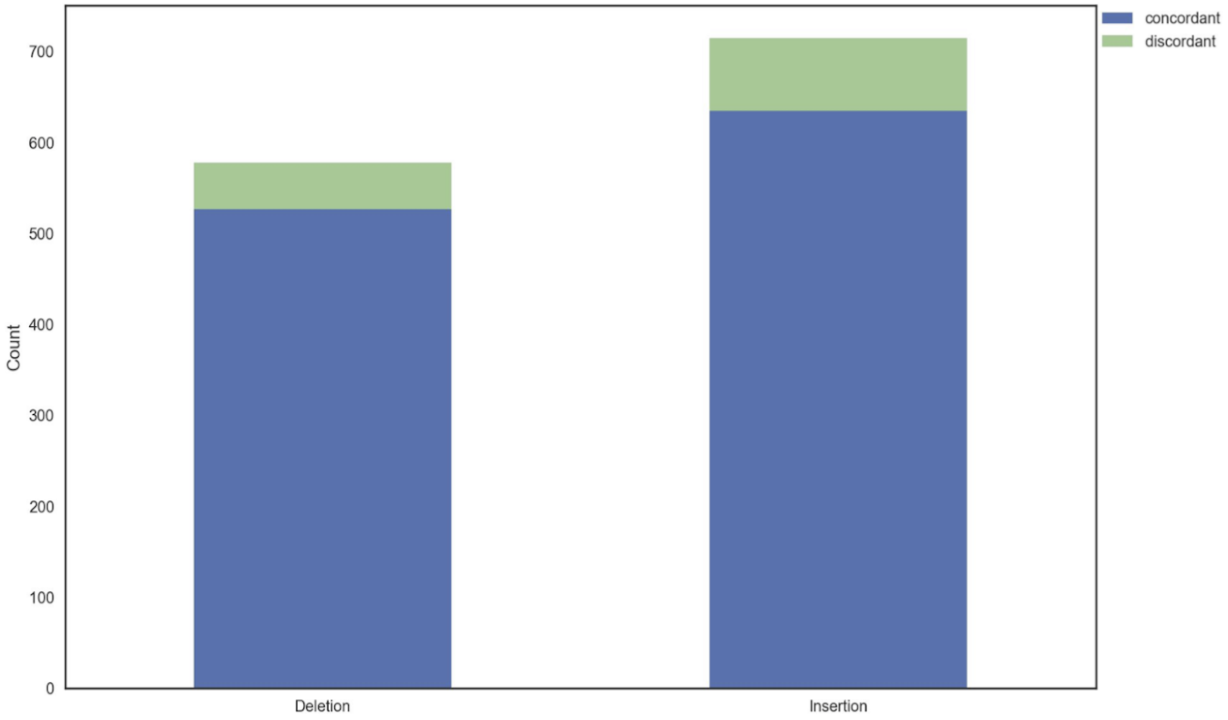


**Supplementary Figure 4.** Evaluation of concordance between Threshold 1 Top Curators (curators that had at least 90.9% concordance with experts) and Threshold 2 Top Curators (curators that had at least 77.7% concordance with experts).

A)

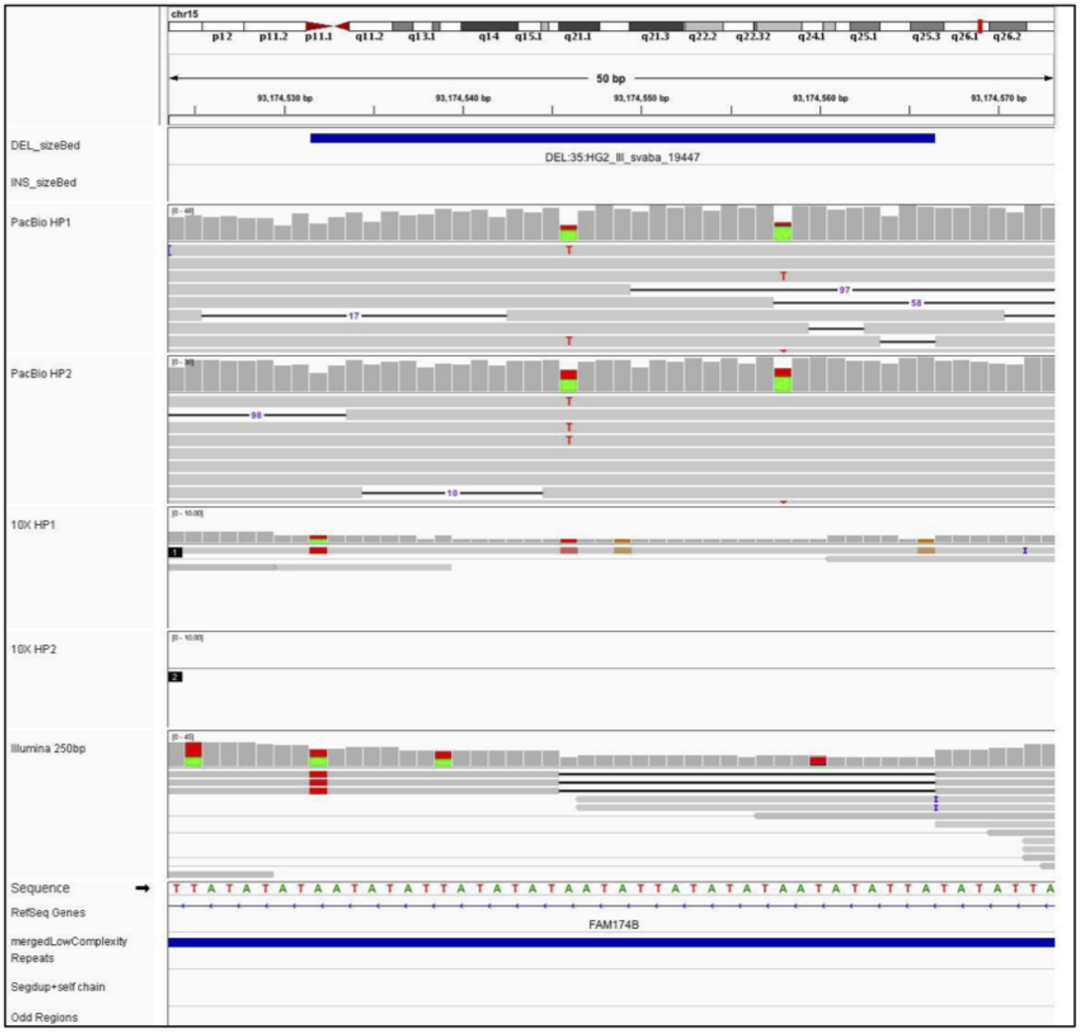
chr15
p12
p11.2
p11.1
q11.2
q13.1
q14
q15.1
q21.1
q21.3
q22.2
q22.32
q24.1
q25.1
q25.3
q26.1
q26.2
50 bp
93,174,530 bp
93,174,540 bp
93,174,550 bp
93,174,560 bp
93,174,570 bp
DEL_sizeBed
DEL:35:HG2_lll_svaba_19447
INS_sizeBed
PacBio HP1
PacBio HP2
10X HP1
10X HP2
Illumina 250bp
Sequence
RefSeq Genes
FAM174B
mergedLowComplexity Repeats
Segdup+self chain
Odd Regions

B)

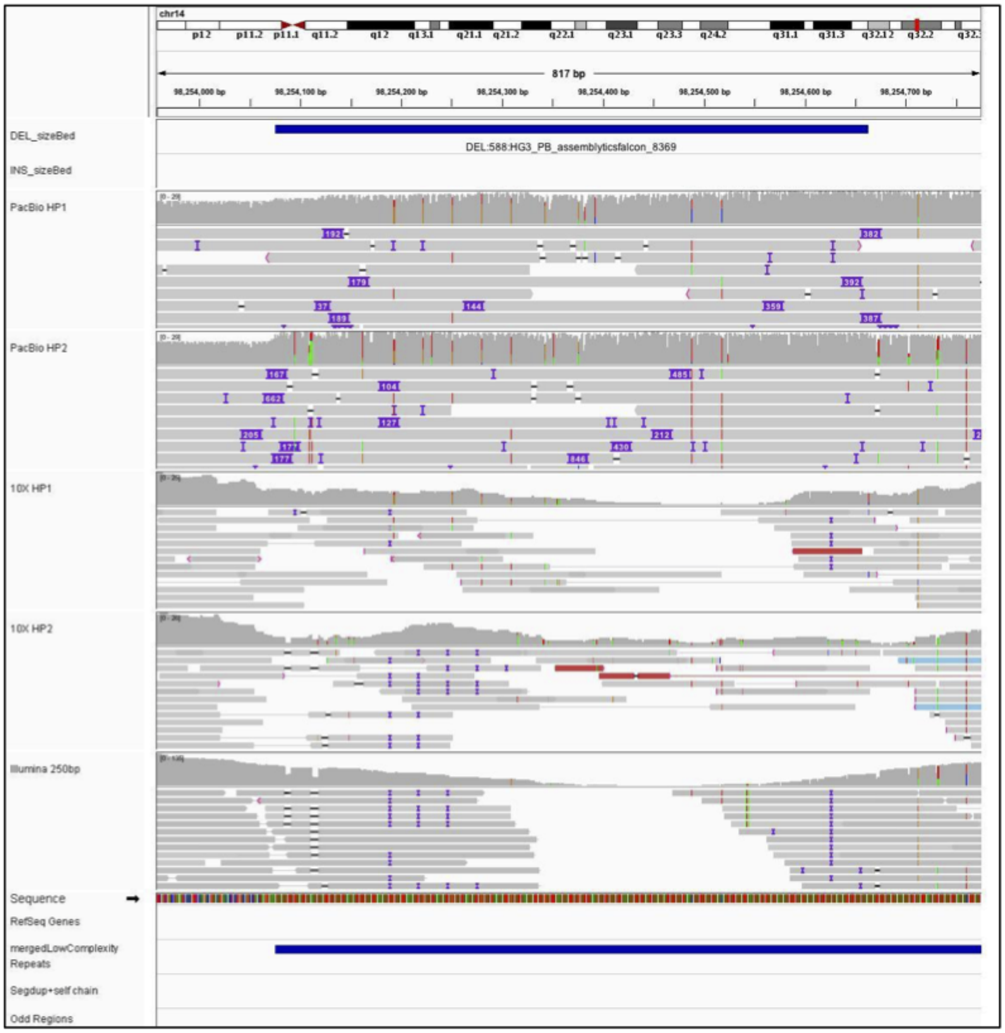


Supplementary Figure **5**. Examples of events that were discordant between consensus labels assigned by curators and the v0.6 high confidence genotypes discordant sites. IGV images showing examples of two events that had less than 50% concordance for the label assigned by the curators. A) small SV call. B) Large SV call.
